# Changes In Ploidy Affect Vascular Allometry And Hydraulic Function In Trees

**DOI:** 10.1101/2021.06.10.447312

**Authors:** M Barceló-Anguiano, NM Holbrook, JI Hormaza, JM Losada

## Abstract

- The enucleated vascular elements of the xylem and the phloem offer an excellent system to test the effect of ploidy on plant function because variation in vascular geometry has a direct influence on transport efficiency. However, evaluations of conduit sizes in polyploid plants have remained elusive, most remarkably in woody species.
- We used a combination of molecular, physiological, and microscopy techniques to model the hydraulic resistance between source and sinks in tetraploid and diploid mango trees.
- Tetraploids exhibited larger chloroplasts, mesophyll cells, and stomatal guard cells, resulting in higher leaf elastic modulus and lower dehydration rates despite the high water potentials of both ploidies in the field. Both the xylem and the phloem displayed a scaling of conduits with ploidy, revealing attenuated hydraulic resistance in tetraploids.
- Conspicuous wall hygroscopic moieties in the cells involved in processes of transpiration and transport advocates a role in volumetric adjustments due to turgor change in polyploids, which, together with the enlargement of organelles, cells, and tissues that are critical for water and photo assimilate transport at long distances, imply major physiological novelties of polyploidy.

## Introduction

Polyploidization and cellular allometry are interrelated processes that pervade the evolutionary trajectory of eukaryotes (Bennett, 1972; Doyle & Coate, 2019). In animal tissues, cell size increases with ploidy, but this is compensated by reducing the number of cells, thus maintaining the final organ size (Kondorosi *et al*., 2000). In contrast, organ size control in plants is more complex, often resulting in a range of sizes across ploidies, despite the consistent cell enlargement associated with ploidy escalations (Stebbins, 1950; Harashima & Schnittger, 2010). Most studies in plant polyploids focus on herbaceous models, concluding that whole genome duplications associate with a general increase in cell volume, but not proportionally in all tissues (Katagiri *et al*., 2016), although the response varies across species. For example, leaf epidermal cells are larger in isogenic tetraploids of *Arabidopsis*, but not in those of *Gossypium* (Snodgrass *et al*., 2017). Comparisons of stomatal sizes across taxa, however, revealed a consistent positive scaling between ploidy and guard cell size (Masterson *et al*., 1994; Beaulieu *et al*., 2008; Simonin & Roddy 2018). Despite the morphological variation associated with ploidy, linking polyploidization with cell function remains challenging, given that an array of factors that affect function arise with whole genome duplication, such as a higher transcript load or the rearrangement of gene expression, arise with whole genome duplication (see Comai, 2005; Iestwaart *et al*., 2017; Jones *et al*., 2017). Unique among plant cells are vascular conduits of the xylem and the phloem, composed of enucleated cells serially connected, whose geometry is determinant for their major function, hydraulic transport. Strikingly, only a handful of studies have related ploidy and vascular cell size, with remarkable exceptions that include the tradeoffs between hydraulics and stress resistance of the xylem in different cytotypes of *Atriplex* (Hao *et al*., 2013), or xylem hydraulic capacitance in *Malus* (Maherali *et al*., 2009; De Baerdemaeker *et al*., 2017). As a result, the correlation between morpho-physiological variation and ploidy remains poorly studied for the xylem, and totally unexplored in the phloem.

Hydraulic transport in vascular plants is governed by the size-dependent hierarchical distribution of conduit elements that form continuous tubes of the xylem and the phloem tissues. Hydraulically conductive cells (tube elements of the xylem or the phloem) vary in size spatially (in an organ dependent fashion), and temporally (during ontogeny), in a manner thought to integrate plant growth and adaptation to environmental conditions. This is particularly relevant in woody plants, which exhibit long life spans, seasonally renovating conduits for maintaining long distance hydraulic transport (Savage, 2019; Savage & Chuine, 2021). Largely, the anatomy and hydraulics of the aerial organs have been studied in isolation, limiting our understanding of the continuous water and photoassimilate transport in woody angiosperms. For example, the structural variation of the elements composing the vasculature is relatively well understood in the trunks, where a universal scaling relationship has been untangled between the geometry of the xylem and phloem conduits and tree height, both in temperate forests or across biomes (Olson *et al*., 2014, 2018; Savage *et al*., 2017; Liu *et al*., 2019). Analogous scaling relationships characterize vessel size in the different vein orders of reticulate leaves (Brodribb & Feild, 2010; Sack & Scoffoni, 2013; Carvalho *et al*., 2018; Gleason *et al*., 2018), but the phloem remains poorly documented, except for recent studies in leaves of some woody plants (Carvalho *et al*., 2017; 2018; Losada & Holbrook, 2019). In reproductive structures, information on the size of vascular conduits is particularly scarce, despite the fact that they constitute major sinks (Zhang *et al*., 2006; Ganino *et al*., 2011). Determining vascular cell size in the continuous tubes that intersect the aerial organs of trees is critical to understand plant productivity and response to internal and external interactions.

Polyploidy may become an advantage for plant confronted with stressful abiotic conditions (van der Peer *et al*., 2020) due to the intrinsic capacity of polyploids to generate physiological novelty (Doyle & Coate, 2019; Roddy *et al*., 2020). Traditionally, agriculture and horticulture have benefited from the use of natural or artificial polyploid genotypes with increased tolerance to stresses such as salinity or drought. Herbaceous polyploid genotypes with these characteristics are present in genera of agronomic interest, such as *Lycopersicum* (Tal & Gardi, 1976), *Raphanus* (Pei *et al*., 2019), or *Fragaria* (Wei *et al*., 2018, 2019), but also in ornamental species such as *Phlox* (Vyas *et al*., 2007), *Chamaenerion* (Maherali *et al*., 2009), or *Saxifraga* (Decanter *et al*.,2020). Comparatively, polyploidy is less frequent in woody species (Ancel Meyers & Levin, 2006), but some results in woody plants point to the capacity of tetraploids to retain more water, such as in seedlings of *Betula* (Li *et al*., 1996), and plants of *Lonicera* (Li *et al*., 2009), or *Populus* (Xu *et al*., 2018). Among fruit tree crops, a handful of polyploids have been studied, revealing sparsely improved physiological properties, such as in the genera *Citrus* (Romero-Aranda *et al*., 1997), *Malus* (Zhang *et al*., 2015; De Baerdemaeker *et al*., 2018), or *Ziziphus* (Wang *et al*., 2019).

The mango (*Mangifera indica* L.) is the fifth most important -in terms of production-perennial fruit crop worldwide (FAO, 2018). Mango was domesticated in tropical Asia about 4000 years ago (Warschefsky & von Wettberg, 2019), but it is increasingly cultivated in subtropical microenvironments of temperate latitudes (Barceló-Anguiano et al., 2021). Most cultivated mango trees are diploid (2n=40) although some polyploids are known (Galán-Saúco *et al*., 2001; García-García *et al*., 2020). A holistic approach to the anatomy and physiology of adult diploid and polyploid trees at the organismal level remains unexplored, not only in mango, but also in most woody perennial species. Here, we take advantage of the availability of autotetraploids of the mango cultivar ‘Kensington’ to compare their anatomical and physiological features with the original diploid cytotype. Our study reveals that, in woody organisms, conduit size increases with ploidy in the aerial organs, most notably in the leaves, implying a first approach to study the effect of genetic dosage on long distance hydraulic transport between organs that are intimately connected through the vasculature.

## Results

### Genotyping and ploidy measurements of mango ‘Kensington’

SSR marker analysis showed no genetic differences between the four trees of the cultivar ‘Kensington’, suggesting that their genotypes were identical. While SSR markers did not offer information on ploidy, flow cytometry analysis confirmed that two of the trees were coincident with the diploid control, and two trees had twice the DNA content, confirming that the latter two trees were autotetraploids and developed from the same diploid genotype.

### Stomata and cell wall chemistry in leaves of diploid and tetraploid mango trees

Cleared leaves revealed that cells of the abaxial epidermis displayed irregular shapes with highly convoluted cell walls, thus projecting only some portions of the cell area toward the abaxial leaf surface (Fig. 1a, b). The stomatal guard cells displayed two well-differentiated zones, the external classic kidney-shaped, and an internal one, devoid of cuticle (Fig. S1), that coincided with the pore aperture, constituting a rounded stomatal complex. Guard cell length was significantly larger in leaves from tetraploid trees (19.1±0.13SE μm long in 4n, vs 14.7±0.10SE μm long in 2n; Fig. 1c), but tetraploid leaves contained a lower stomatal density (410±6.0SE stomata per mm^2^) compared with diploids (715±10.0SE stomata per mm^2^).

**Fig. 1.**
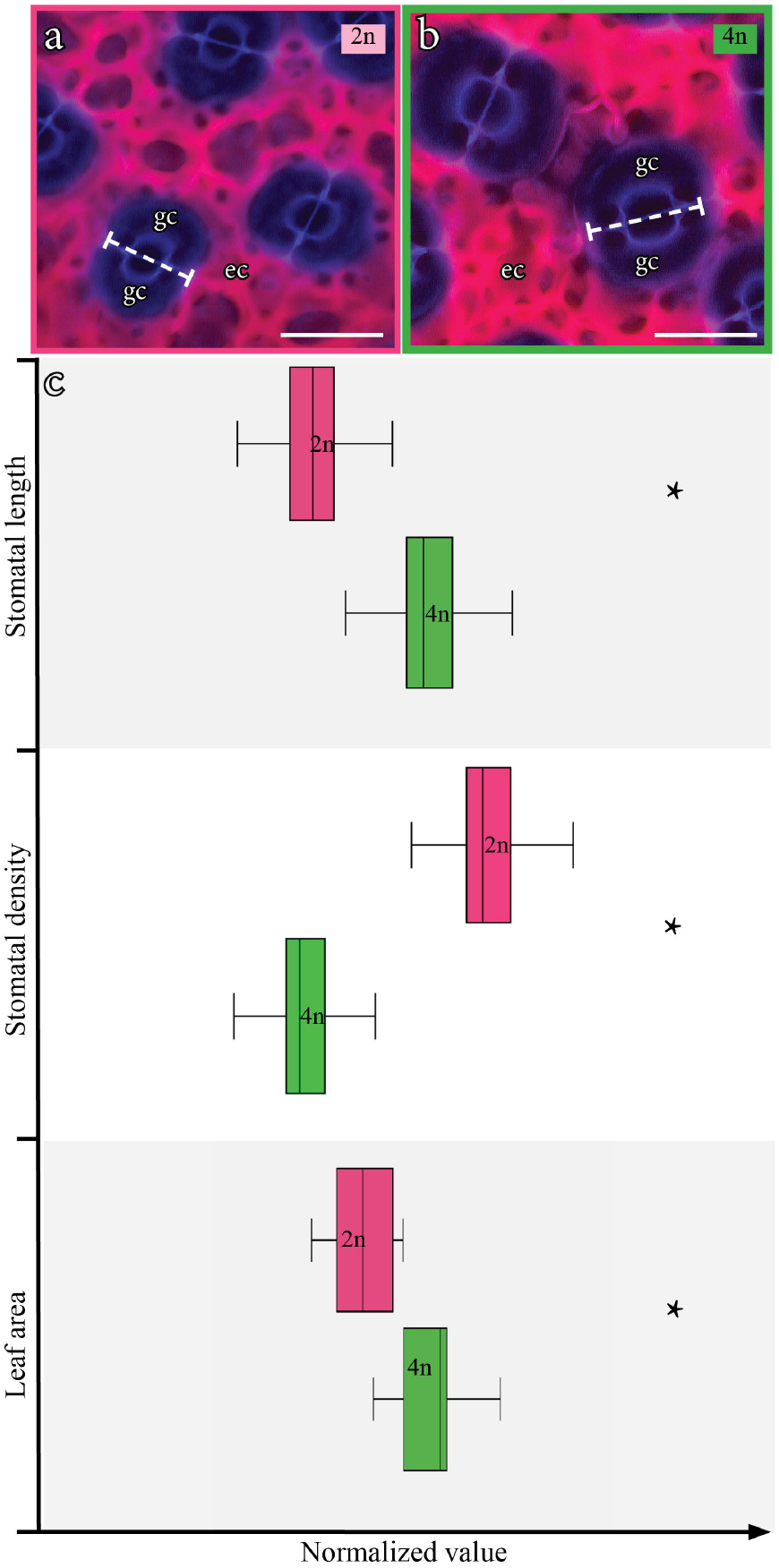
Stomatal morphology and metrics in diploid (pink) and tetraploid (green) mango leaves. **(a)** Stomata of diploid mango leaves after staining and clearing: the purple guard cells (gc) show two differentiated areas, the central pore, devoid of cuticle, and the kidney shaped wider parts; pink color displays the convoluted cell walls of the epidermal cells (ec) projecting toward the abaxial leaf surface. **(b)** Stomata of tetraploid mango leaves. **(c)** Values of stomatal length (quantified as noted by the dashed lines), density and leaf area in diploid (pink), and tetraploid (green) mango leaves normalized to their maximum value for comparison; asterisks reveal significant differences between ploidies at *P* ≤ 0.001. **(a-b)**, 3D maximum projections of whole mounts from cleared leaves of mango, stained with Feulgen reagent, and imaged from the abaxial side. Scale bars = 20μm.

### Water potential (□) and leaf water content in diploid and tetraploid mango trees

Measurements of leaf and stem water potential at midday showed high value ranges during winter (January and February), and at the beginning of the spring season (March) (−0.2 to –0.5MPa), without differences between individuals or ploidy levels (Fig. S2). Despite the fact that the trees were irrigated to full capacity all year round, both stem and leaf water potential of the tetraploid trees were significantly lower during the summer season, when temperatures were higher and no rainfall took place.

Leaf mass per area was significantly higher in tetraploid (187.28±6.7SE g·m^-2^) than in diploid (152.83±5.0SE g·m^−2^) trees (n=24). Despite the fact that tetraploids had a higher water fraction per area (245.65±5.4SE g·m^−2^ tetraploids, and 191.95±4.2SE g·m^−2^ diploids n=42), diploids lost a higher percentage of water through time after bench-dry dehydration (Fig. 2a). Small changes in the relative water content correlated with a sharp drop in the leaf water potential in the leaves of both ploidies (Fig. 2b). Yet, tetraploid leaves reached turgor loss at higher relative water content, resulting in a higher elastic modulus at turgor loss (i.e. more rigid), and higher apoplastic fraction of water after losing turgidity (*P*<0.001, Table I).

**Fig. 2.**
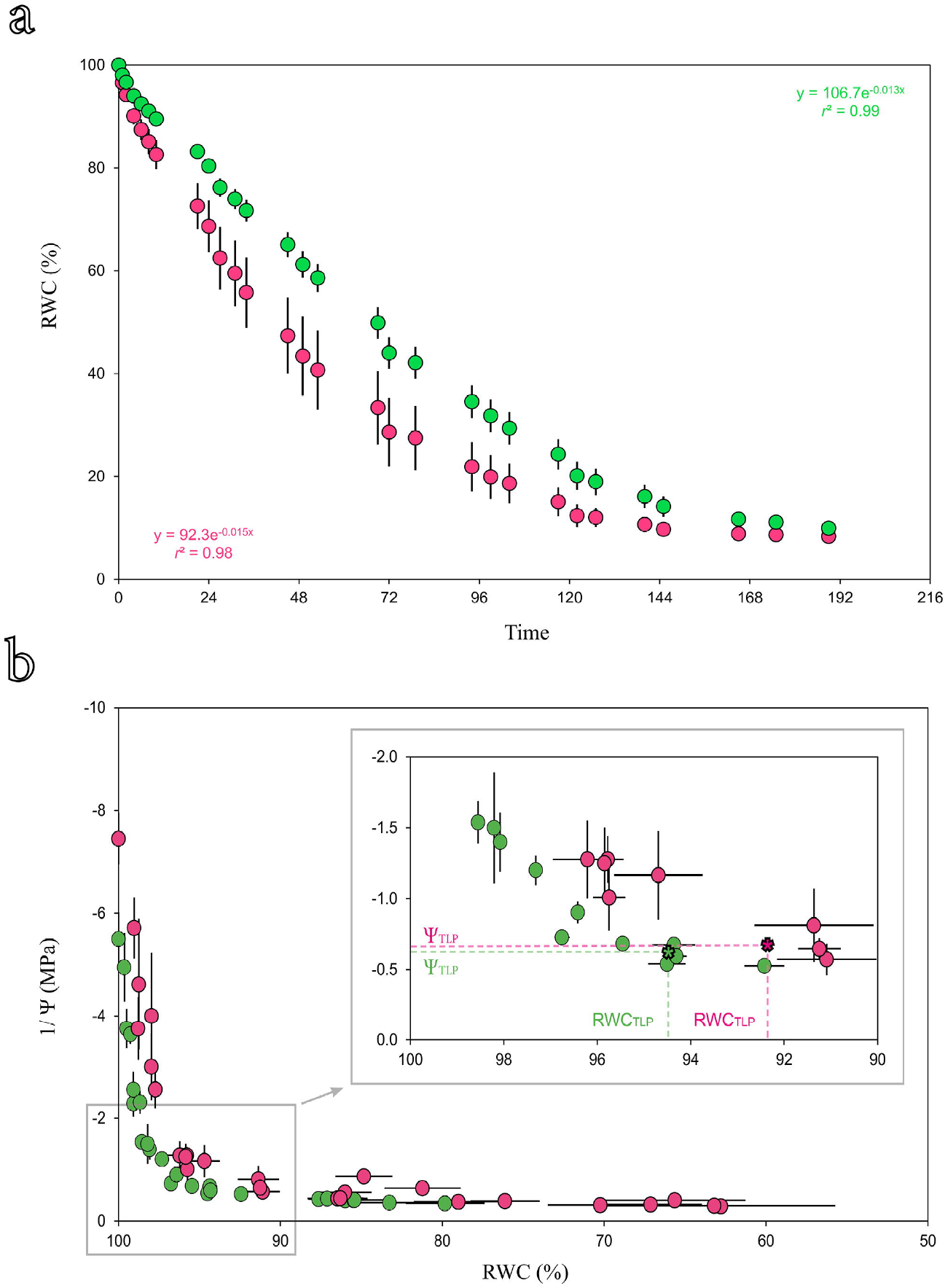
(a) Loss of relative water content (RWC) with time (in hours) of diploid (pink) and tetraploid (green) mango leaves. (b) Pressure-volume (P-V) curves of diploid (pink) and tetraploid (green) mango trees. Vertical and horizontal bars represent the standard error each axis at a *P* ≤ 0.05; inset (grey square in B) shows a magnified view of the plot area where the turgor loss point (TLP, asterisks) is extrapolated for each ploidy.

**Table I.**
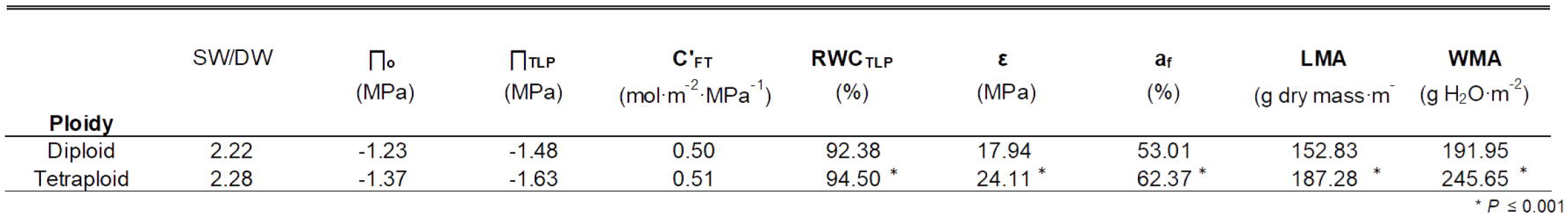
Extrapolated parameters from the P-V curves, leaf and water mass areas. Relationship between saturated weight and dry weight (SW/DW); osmotic potential at full turgor (Π_o_) and at turgor loss point (Π_TLP_); capacitance at full turgor (C’_FT_); relative water content at turgor loss point (RWC_TLP_), elastic modulus (□), apoplastic water fraction (a_f_), leaf mass area (LMA), and water mass area (WMA). Asterisks represent significant differences between ploidies at either a *P* ≤ 0.001 (*).

### Leaf anatomy of diploid and tetraploid mango trees

Cross sections unveiled the distribution of different foliar tissue layers (Fig. 3a). The adaxial epidermis was composed of squared cells, and, underneath, a columnar layer of packed palisade cells. Abaxially, the rounded cells of the spongy mesophyll (Fig. 3b) were interspersed with vascular bundles, and were larger in the tetraploid than in the diploid leaves (Fig. 3c). Similarly, the plastids were also larger in the tetraploid leaves compared with the diploids, as shown by the immunolocalization of the branched pectin epitope recognized by the LM26 monoclonal antibody (Fig. 3b, c). A high concentration of this epitope was massively exposed between the cell membrane and the wall in cells of the abaxial epidermis, reflecting their convoluted perimeter (Fig. 3d, e).

**Fig. 3.**
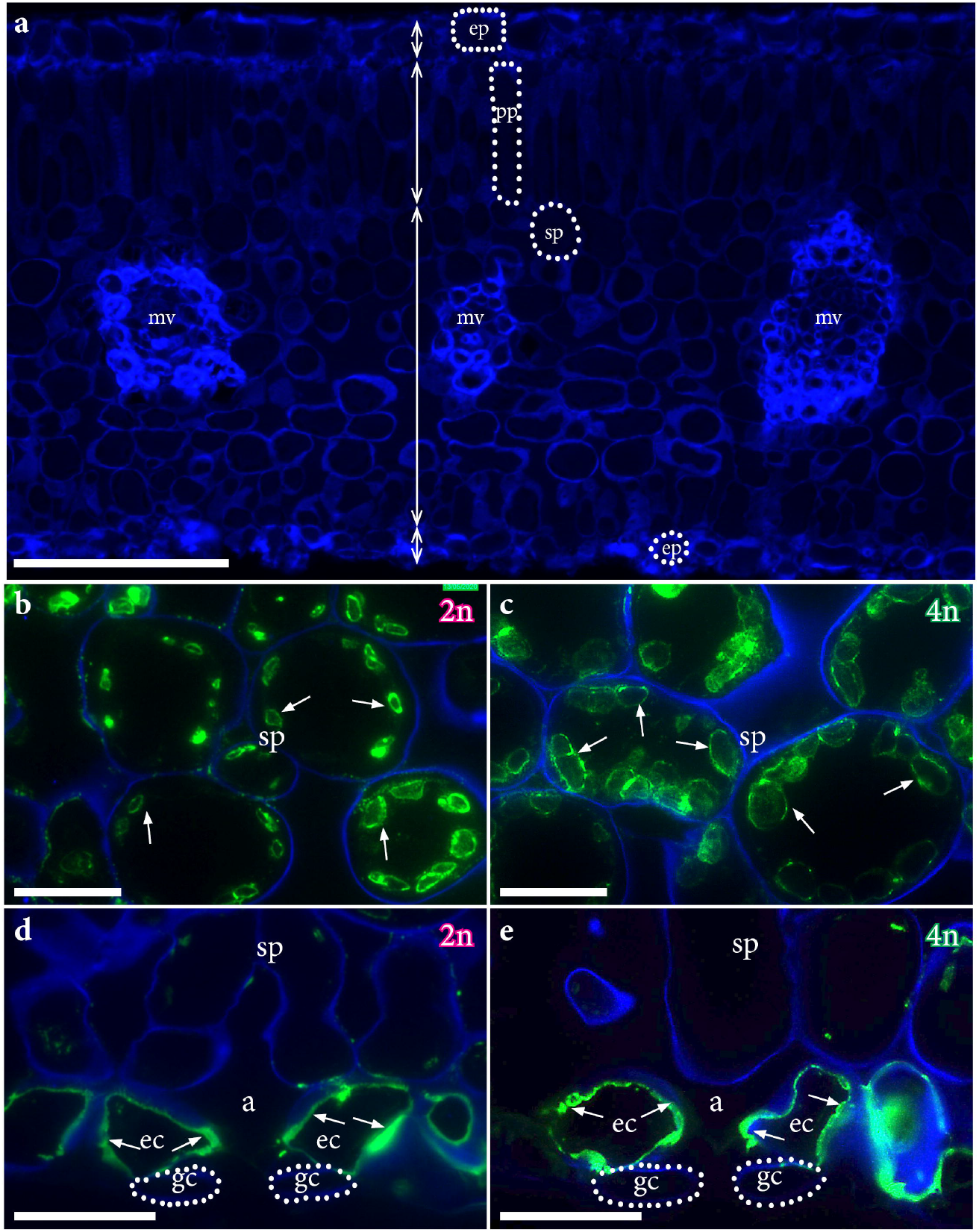
Anatomy of mango leaves and immunolocalization of branched pectins. **(a)** Cross section of a mango leaf displaying the different tissue layers from the adaxial (top) to the abaxial surface (bottom), delimited with double headed arrows. Dotted lines indicate the cell profiles in each tissue type. **(b)** Close-up of the spongy mesophyll with the plastid membrane stained in green (arrows) indicating the presence of a branched pectin epitope recognized with the LM26 monoclonal antibody in diploid leaves. **(c)** The plastids in the spongy mesophyll cells of the tetraploid leaves further localized the epitope, revealing their larger size. **(d)** Abaxial leaf side showing the presence of the epitope pervading the membrane of the epidermal cells, and correlate with the convoluted character observed in cleared leaves from Figure 2 (arrows). **(e)** similar labelling in tetraploid leaves (arrows). A, aereal; ec, epidermal cell; ep, epidermis; mv, minor vein; pp, palisade parenchyma; sp, spongy parenchyma. Scale bars: **(a)** = 100μm; **(b-e)** = 20μm.

### Vascular element geometry in leaves, stems and inflorescences of diploid and tetraploid mango trees

Evaluations of the conduit elements composing the xylem displayed a quite constant length across organs, varying mostly their radii (Fig. 4a,c,e). In turn, phloem conduits varied in both length and radii across organs (Fig. 4b,d,f). Within the same areas of the stems, the length of the vessel elements was not much different to the length of the sieve tube elements, although the sieve cells were five times thinner than the vessel cells (Fig. 4a,b). Strikingly, vessel elements were only twice as wide than sieve tubes in both inflorescences (Fig. 4c,d) and leaves (Fig. 4e,f).

**Fig. 4.**
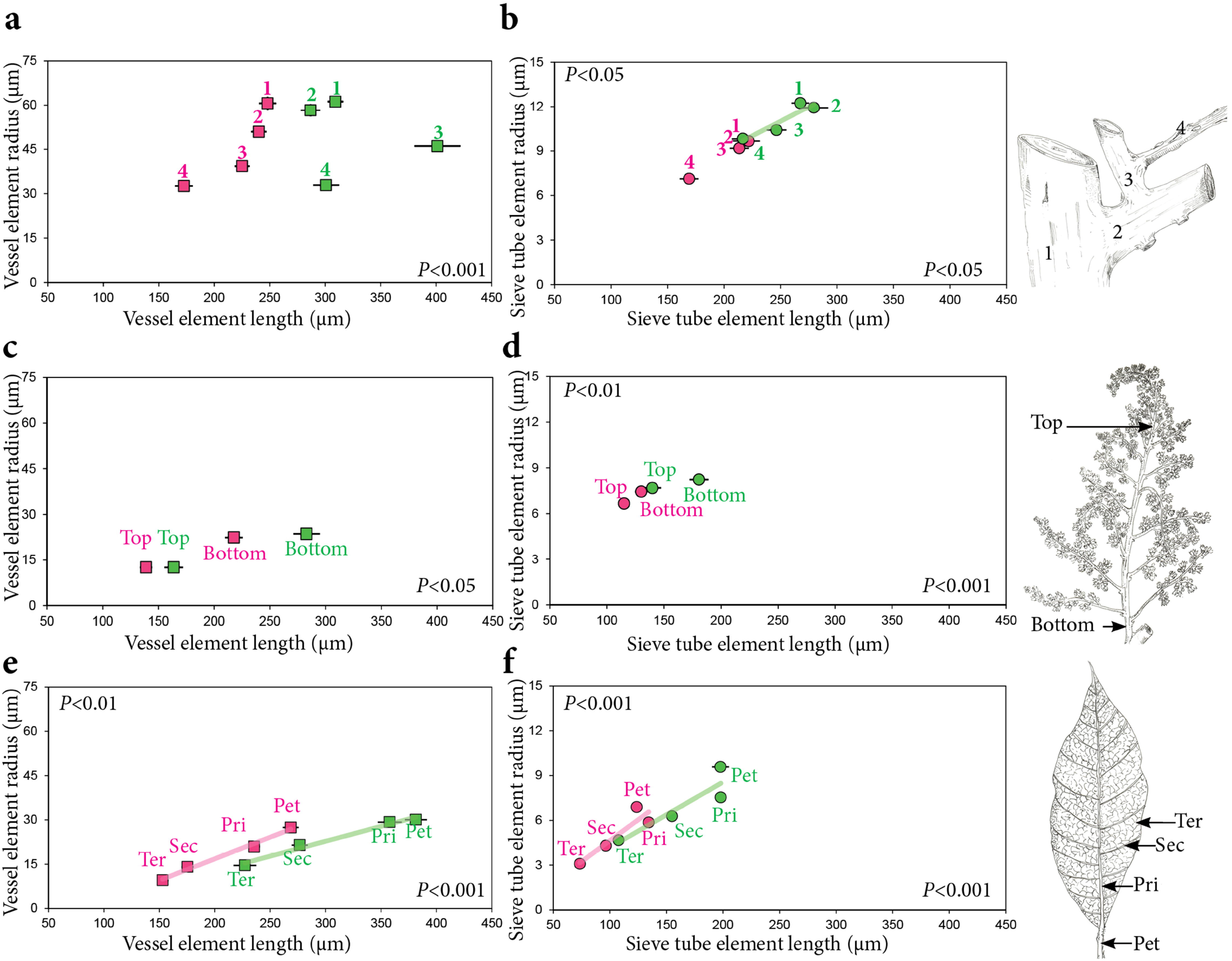
Geometry of the vascular elements at different organs in diploid (pink) and tetraploid (green) mango trees. Left panels show the correlation of radius and length in the vessel elements of the xylem in the inflorescences **(a)**, the stems **(c)**, and the vein orders of leaves **(e)**. Right panels display the correlation between radius and length of the sieve tube elements of the phloem in the same localizations. Pairwise comparisons of the length or radius of the vascular elements displayed significant differences between ploidies at the specified *P* value in each axis at similar locations. Artwork kindly provided by Laura Carrera.

Comparisons between ploidies revealed that, in the branches, the radius of the vessels was similar between ploidies, but the tetraploids displayed longer vessel cells than the diploids (*P*<0.001; Fig. 4a). The tetraploid sieve tube elements of the phloem in stems were longer (*P*<0.001) and wider (*P*<0.05) than those of the diploids in all branch diameters (Fig. 4d). In the inflorescences, the vessel elements at the bottom of the rachis were longer and wider than those at the tip in both ploidies (Fig. 4a), and they were significantly longer –but not wider– in the tetraploids than in the diploids (*P*<0.05). The sieve tube elements of the tetraploid inflorescences were significantly larger (*P*<0.001) and wider (*P*<0.05) at both locations of the rachis (Fig. 4b). In the leaves, the vessel elements, which scaled linearly with the vein order, were longer and wider in tetraploids (Fig. 4e). Similarly, the tetraploid sieve conduit elements were longer and wider than those of the diploids (*P*<0.001; Fig. 4f).

The simple perforation plates of the vessels were circular or elliptical at all locations along the tree, and their diameter was, in general, 10% smaller than that of the vessel elements. In contrast, the sieve plates of the phloem were composed of several areas (multiple connections between tubes, Fig. 5a, d). These individual sieve areas enlarged linearly from the tertiary vein order to the widest stem (Fig. S3), and accordingly, the pore size followed a similar pattern (Fig. 5b, c, e, f). However, while no consistent differences were observed in pore size between ploidies (Fig. 5g), the tetraploid sieve tube elements contained a higher number of sieve areas per end plate, resulting in a higher total estimated number of pores at all phloem locations than those of the diploids (Fig. S4).

**Fig. 5.**
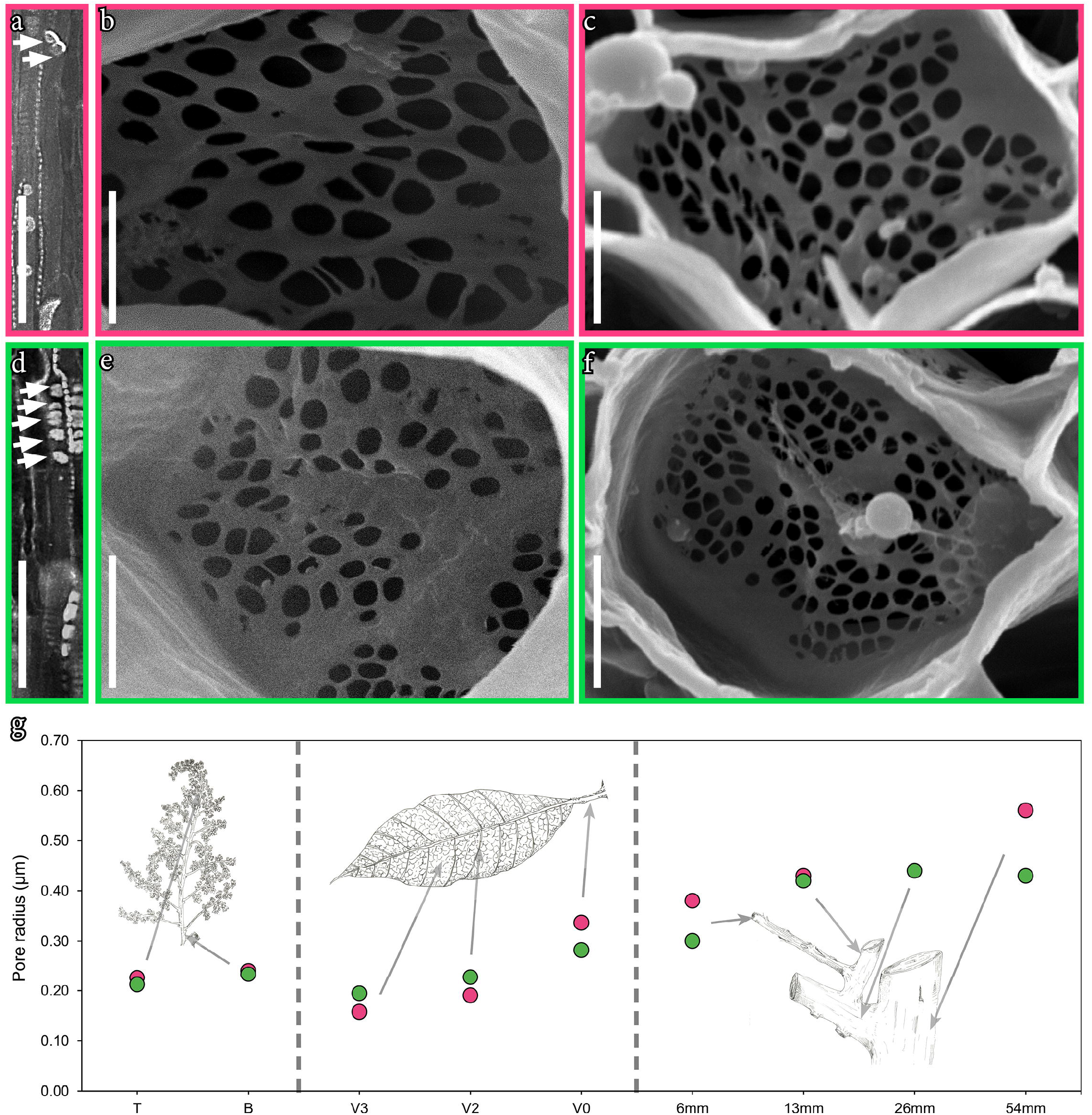
Pore radii across the phloem pathway of diploid (pink), and tetraploid (green) mango trees. **(a)** Anatomy of the sieve tube element of diploid mango trees, showing compound plate connections (arrows) per end tube. **(b)** Sieve area in the stem of a diploid mango tree. **(c)** Sieve area in the petiole of diploid mango leaves. **(d)** Sieve tube of a tetraploid mango tree, with more sieve areas composing the plate connections than in diploid mango trees (arrows). **(e)** Sieve area of the stem from a tetraploid mango tree. **(f)** Sieve area of a petiole of tetraploid mango leaves. **(g)** Radii of pores in the sieve plates comparing diploid (pink) and tetraploid (green) mango trees. X axis denotes the location in the inflorescence (T, top; B, bottom), vein order (V3, terciary veins; V2, secondary veins; V0, petiole), or stem diameter (6mm, 13mm, 26mm, 54mm). Bars (not visible in the plot) represent the standard error at a *P* ≤ 0.05. A, D. Fluorescence images of the sieve tubes stained with aniline blue for callose and with calcofluor white for cellulose in the cell walls. B,C,E,F. Scanning electron microscope images of the individual sieve areas forming the compound sieve plate (circles). **(a,d)** scale bars = 50μm; **(b,c,e,f)** scale bars = 5μm.

### Hydraulics of the xylem and the phloem in diploid and tetraploid mango trees

Sap is transported in opposite directions -and pressures-through the xylem and the phloem (Fig. 6a), but the lumen resistance per length of both vascular tissues decreased exponentially from the minor veins of the leaves to the base of the stems (Fig. 6b). Due to the thinner conduits of the phloem, the lumen resistance was two to four orders of magnitude higher in that tissue than in the xylem. The lumen resistance of the vessel tube per distance decreased by four orders of magnitude toward the base of the stems, following a power law but this decrease was sharper in the diploids (Fig. 6b). While the lumen resistance of the inflorescence rachis was similar to that of the current year branches, where they typically develop, the lumen resistance of the vessel tubes increased sharply toward the inflorescences. Comparison between ploidies revealed a lower lumen resistance of both the sieve and the vessel tubes in the tetraploids, contributed mainly by the wider conduit size, which offset the resistance added by longer conduits of tetraploids.

**Fig. 6.**
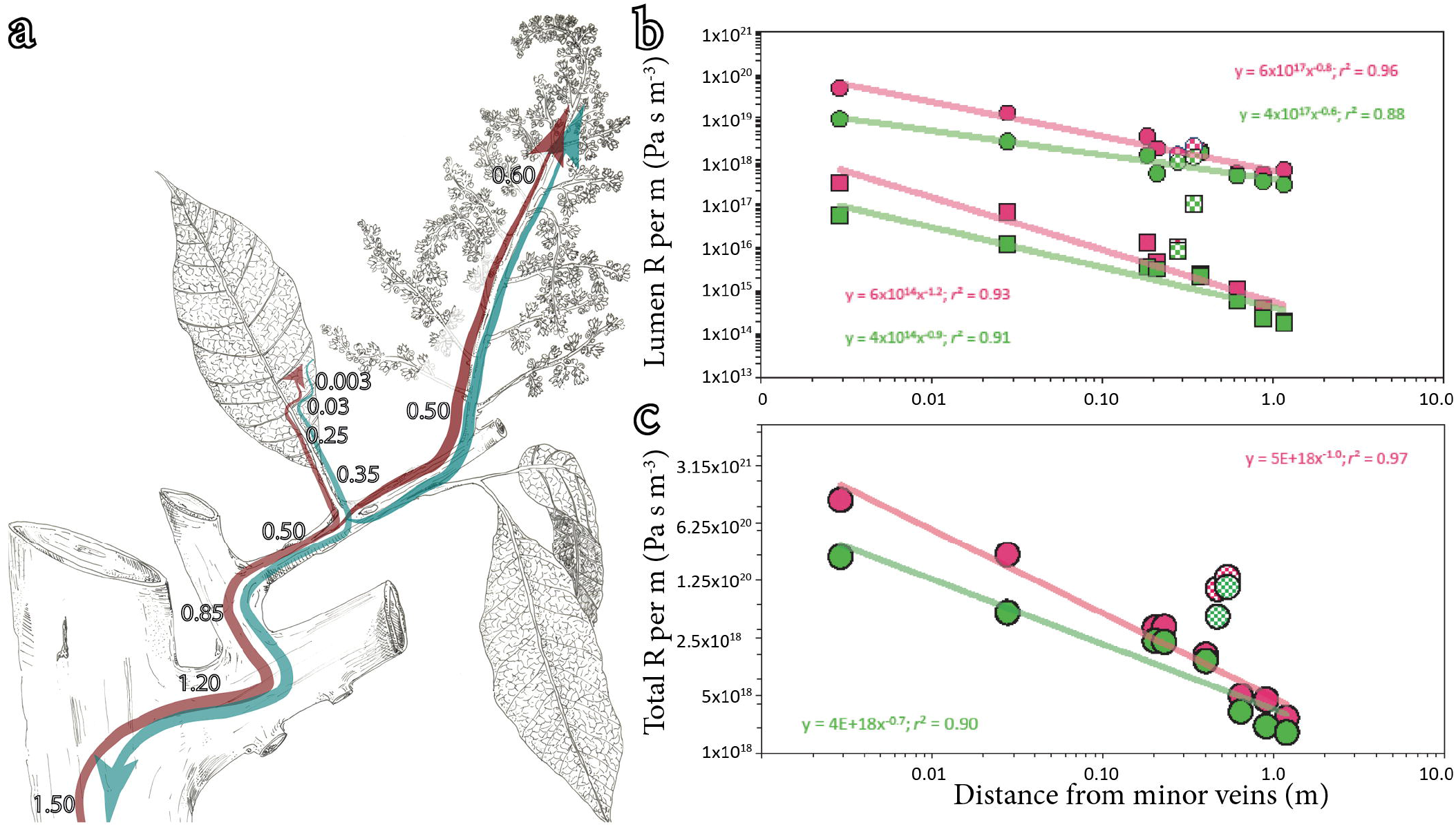
Axial resistance of the vascular elements of the xylem and the phloem along the transport pathway of diploid (pink) and tetraploid (green) mango trees. **(a)** Continuous pathway of the xylem (magenta), and the phloem (turquoise) with arrows representing the directionality of the sap flow within each vascular tissue; numbers represent the cumulative distance from the minor veins in m; note the secondary unidirectional pathway toward the upright inflorescences. **(b)** The lumen resistance per m of the sieve tube (circles) was two to four orders of magnitude higher than the vessel tube (squares) per m, but both decreased exponentially with distance from the minor veins. The tetraploids displayed lower resistances that diploids in both the xylem and the phloem tubes, particularly in the leaves; while the lumen resistance per m of the phloem was maintained in the inflorescences, the vessel resistance increased sharply toward the inflorescences (hatched symbols). **(c)** Total resistance per m of the sieve tube decreased exponentially with distance from the sources (i.e. minor veins) for the vegetative organs, but the reproductive organs (hatched circles) increased the total resistance due to the tiny pore size of the sieve plates. The tetraploid tubes tend to have lower resistances than the diploids, especially in the leaves. Exponential regressions at a *p*<0.05.

The major effect of pore number in the total resistance of the sieve tube of the phloem was evidenced by a lower total resistance -per length-of the tetraploids (Fig. 6c). The total resistance scaled down exponentially in the vegetative organs (from leaves to stems), with tetraploids displaying attenuated hydraulic resistances compared with diploids. Strikingly, the tiny pore sizes of the inflorescence rachis implied a significant increase in the total sieve tube resistance toward the reproductive tissues (Fig. 8c), although the number of pores evaluated was small due to the abundant latex in these organs, which blocked observation of most sieve plates.

## Discussion

### Enhanced leaf hydration and water retention in tetraploids

Our evaluations revealed that tetraploids displayed a higher proportion of dry mass per leaf area, which is consistent with their larger cells of the spongy mesophyll. Previous reports correlated dry mass and ploidy in colchicine-induced herbaceous polyploids (Decanter *et al*., 2020 and references therein), and in some woody polyploids, such as *Populus* (Zhang *et al*., 2019) or *Lonicera* (Li *et al*., 2009). We further noticed a higher fraction of water per area unit in tetraploids, but leaf bench dehydration rates were slower than diploids, pointing to lower epidermal conductance of tetraploids. Under field conditions, diploid and tetraploid mango trees displayed similar midday leaf and stem water potentials under field conditions (between −0.10 and −0.50MPa), but when leaves were cut, tetraploids reached turgor loss point at higher relative water content than diploid leaves, resulting in higher values of apoplastic water. Recent compilations comparing trees growing in different environments reported variability of this parameter among trees, fluctuating from 15% to 80% (Nadal *et al*., 2019). While our values in mango fall within ranges reported for most studied tropical evergreen trees, these unusual high levels of water in the apoplast could likely associate with the pervasive laticiferous canals embedded in the phloem tissue along the leaf veins (Barceló-Anguiano et al., 2021), which are typically filled with aqueous compounds. Additionally, tetraploid leaves showed a higher elastic modulus, a parameter that describes the relationship between turgor and cell volume and reflects foliar cell rigidity (Barlett *et al*., 2012; Trifilo *et al*., 2016; Nadal *et al*., 2019). If stomatal closure is linked to the turgor loss point, a higher elastic modulus may be useful during periods of water scarcity, extending the time in which water is retained in the leaves compared with diploids, as observed, for example, in polyploid rootstocks (for a review see Ruiz *et al*., 2020). Pressure-volume relationships have been extensively used in the study of leaf and stem hydraulic properties of woody species (Sack *et al*. 2003; Nadal *et al*., 2019), but rarely comparing conspecific individuals with different ploidies, such as in seedlings of *Betula*, which also exhibited a higher water content at turgor loss for tetraploids (Li *et al*., 1996).

### Foliar cell allometry in isogenic polyploids

A higher water fraction compartmentalized in tetraploid leaves correlates with the morphology of leaf internal tissues. For example, bigger mesophyll cells of tetraploids create larger surfaces, and may thus expose a higher area of cell-wall related hygroscopic materials that retaine water. Enlarged mesophyll cells in polyploids were previously observed in a handful of trees, such as *Citrus* (Romero-Aranda *et al*., 1997). Strikingly, localization of a branched pectin in the plastids revealed that tetraploid plastids were twice as large as diploid plastids, evidencing size-related effects of ploidy on key organelles for photosynthesis efficiency, and, thus, plant performance (discussed by Doyle & Coate, 2019). In addition, the total number and size of plastids appear as a key feature for the mesophyll conductance, recently stated as a neglected parameter with importance to improve the physiology of angiosperms (Nadal & Xiong, 2020).

Our quantifications showed longer but less dense stomatal guard cells in the tetraploids, consistent with the pattern observed across polyploids in other angiosperms (Beaulieu *et al*., 2008). Lower stomatal densities have been recently reported as better predictors of drought tolerance than guard cell length in herbaceous crops such as barley or rice (Harrison *et al*., 2019; Caine *et al*., 2018; Hughes *et al*., 2017). Both density and size of the guard cells influence the degree of transpiration, a process that requires volumetric adjustments of the neighboring epidermal cells, typically overlooked in most studies. We revealed for the first time in an angiosperm the presence of a specific branched pectin in the plasma membrane of the cells of the abaxial epidermis of mango leaves. These cells display highly convoluted walls, suggesting a role on the turgor-mediated volumetric changes taking place in the abaxial epidermis during the stomatal cycle of transpiration. Yet, additional work will determine the ubiquity of these pectins in the abaxial epidermis of other angiosperm leaves.

### Axial allometry and the effect of ploidy on xylem hydraulics

Our work reveals that, compared to the diploid trees, the vessels of tetraploid trees were significantly longer at all axial locations in the plant, but particularly longer and wider through leaf veins. Comparisons of vessel element size along the transport pathway in isogenic lines has remained elusive so far, but previous works pointed to a direct correlation between ploidy and hydraulics of the xylem in the stems of *Atriplex*, confirming that polyploids of this genus colonize drier habitats due to an enhanced drought tolerance (Hao *et al*., 2013). A more recent work with apple trees suggests a higher hydraulic capacitance of tetraploid stems, which correlates with a higher vessel area compared with diploids (Baerdemaker *et al*., 2018). Although such studies may explain a buffering capacity of tetraploids during dry periods, most information on drought tolerance of tetraploids comes mostly from herbaceous crops such as tomato (Smiths, 1946; Tal & Gardi, 1976; Lewis, 1980; Lumaneret, 1987). Our correlations between vessel element size and ploidy in mango leaves suggest that tetraploids displayed lower hydraulic resistances to water transport than the diploids. As a result, under water deficit, the diploids may require higher xylem tensions than tetraploids to support the same transpiration rates, but this needs verification *in vivo*.

We further confirmed a hierarchical distribution of vessel elements along the xylem tubes in mango, narrowing acropetally along the transport pathway, from the major trunk through the thinnest branches, the petiole and leaf veins, in line with the cohesion-tension theory (Dixon & July, 1895). This architectural continuity follows the scaling observed in the trunks of several temperate forest trees (Savage *et al*., 2017; Olson *et al*., 2018; Liu *et al*., 2019). Our calculations show that the increased resistance is of four orders of magnitude from the base of the stems, through the tapering branches, to the minor veins of the leaves. Mango leaves display a linear scaling of the vessel geometry (length and radius) across vein orders, similar to previous evaluations of xylem architecture in angiosperm leaves with reticulated venation (Feild and Brodribb, 2013; Carvalho *et al*., 2017). Together, these results support the hydraulic segmentation hypothesis, which states that leaves exhibit the highest xylem hydraulic resistance (Zimmerman, 1983; Sack & Holbrook, 2006). Our work adds further evidence on the hydraulic segmentation of the reproductive inflorescences, which, in mango, displayed a xylem hydraulic resistance comparable to values of the secondary veins of leaves due to their constricted geometry. While the segmented profile of the xylem toward inflorescences may serve as a safety margin against cavitation (Levionnos *et al*., 2020), this architecture implies that the inflorescences are under relatively high xylem tensions. Strikingly, the ratio vessel radii: sieve tube radii in the inflorescences and leaves is 1:2, much smaller than the 1:5 of stems, evidencing organ-dependent cambial activity that reflects in the conducting balances between the xylem and the phloem, poorly studied in reproductive organs (Ganino *et al*., 2011).

### Relaxed phloem hydraulic resistance in polyploids

The sieve tube elements of the phloem in tetraploid trees had larger geometries than the diploid ones at all locations along the transport pathway in mango trees. To our knowledge, this is the first report comparing the architecture of the phloem between isogenic individuals with different ploidies. These results strongly suggest that the geometrical parameters of the sieve tube elements scale with ploidy, and, consequently, the resistance of sieve tubes is lower in the tetraploids.

Despite the general volumetric increment of the phloem conduits with ploidy, not all the tube features varied coordinately. For example, there are no differences in pore size or sieve area size between ploidies. Pore size is the most influential constraint on sieve tube conductivity, affecting this property as to the fourth power (Jensen *et al*., 2012; Knoblauch *et al*., 2016). Conversely, the number of sieve areas per compound sieve plate (the number of connections per end tube) was significantly higher in the tetraploids at all locations of the transport pathway, affecting the overall tube resistance. Indeed, our calculated phloem hydraulic resistance, which is in line with previous works with species from different phylogenetic backgrounds (Losada & Holbrook, 2019; Carvalho *et al*., 2018; Clerx *et al*., 2020), emphasize the fact that comparisons between isogenic lines with different ploidies are key to understand the relationship between gene duplication and vascular novelty in individual taxa.

Relaxation of the phloem hydraulic resistance in tetraploid leaves has important implications for long distance photoassimilate transport, especially during periods of water scarcity (Liesche, 2020). Theoretical frameworks argue that under extreme drought, the viscosity of the phloem sap increases up to a threshold above which the transport is impaired (see Sevanto, 2018). Predictably, during phloem loading, the wider tubes of tetraploids would need higher amount of solutes to reach a similar pressure compared with those of diploids. While this may imply a higher fraction of water uptake from the xylem, tetraploids would reach the maximum viscosity later than in diploid sieve tubes under the same drought conditions, conferring to tetraploids an expanded window of phloem vulnerability prior to collapse. Empirical evidence to test this hypothesis is so far lacking.

Our results are in line with an intraorganismal scaling relationship of the phloem from source to sink, forming a continuum leaf-branch-inflorescence or leaf-branch-base of the trunk. The scaling geometry of the sieve tube elements was consistent in all studied organs, including the leaves, the stems and the inflorescences. We provide more evidence that adds to recent works showing a correlation between the height of the tree or the branch diameter with sieve tube structure (Savage et al., 2017; Losada & Holbrook, 2019; Clerx *et al*., 2020; Barceló-Anguiano et al., 2021). While our measured pore sizes in the stems fall within the low range of those previously reported in several temperate trees (Liesche *et al*., 2017; Savage *et al*., 2017), the pore size of the sieve tube elements in leaves and inflorescences remained unexplored so far. Our study examined for the first time in an angiosperm the scaling of pores associated to sieve tubes in both the leaves and the inflorescences. In the leaves, the higher pressures required for transport are in agreement with the Münch hypothesis of phloem transport (Münch, 1930), by which movement in the phloem results from osmotically produced pressure gradients between source and sink organs. In the inflorescences, however, we reveal a significant increase of sieve tube resistance, which appears counterintuitive according to this theory. We predict that the pressure gradient into reproductive sinks could be large since they are growing rapidly and thus could lower the phloem pressure. Photo assimilate unloading during the flowering period could also be actively regulated by osmolites, provided by the massive laticiferous canals in the inflorescences (Pickard, 2007).

## Conclusions

Polyploidization in plants has been widely studied in angiosperms from an evolutionary perspective, but rarely from the angle of hydraulics. Our evaluations with genetically identical mango trees with duplicated chromosome cargo revealed a strong correlation between ploidy and the size of key cells involved in plant physiology, such as the stomata, the mesophyll cells, and the conduits composing the xylem and the phloem vascular tissues. These anatomical innovations, along with the presence of flexible-related compounds in the walls of cells that are directly or indirectly involved in essential physiological functions, such as the abaxial epidermis or the sieve tube elements, point to an enhanced physiological performance of the tetraploids. Autotetraploid resilience needs may rely on the capacity to build stiffer leaves that dehydrate at slower rates, opening new avenues for future investigations on drought resistance in trees.

## Experimental procedures

### Plant material

Mango trees (*Mangifera indica*) of the cultivar ‘Kensington’ located in the subtropical microenvironment of Málaga province in southern Spain, maintained at the mango germplasm collection of the IHSM La Mayora (4□ 2’ 24.32’’ longitude and 36□ 45’ 32.75 latitude), were used. We evaluated four 6-year-old well-irrigated trees (two diploid and two tetraploid), grafted onto the diploid clonal rootstock cultivar ‘Gomera4’. The trees are maintained with the canopy about two meters aboveground. At the reproductive phase, they produce apical inflorescences positioned upright (Fig. S5a) with numerous male and hermaphrodite flowers per inflorescence (Fig. S5b). During the initial fructification, massive abscission of flowers leaves an average of 1 to 2 fruits per inflorescence (Fig. S5c), normally located in the apical part (Fig. S5d).

For sampling, each tree was divided into three main areas: leaves, in which the major veins (petioles, primary, secondary, and third order veins) were collected, inflorescences, in which the bottom and top part of the main rachis were sampled, and stems, in which four diameter ranges (5-7mm, 13-15mm, 23-25mm, and 43-46mm) were selected. We sampled three leaves from each ploidy (n=12), three inflorescences per tree (n=12) and three branches oriented to all directions in each diameter range (n=12).

### Flow cytometry and genotyping

Young leaves were chopped (0.5 cm^2^ pieces) into a buffer solution designed for nuclei extraction (Cystin UV Precise T Kit), filtered with a nylon filter of 30 μm pore size, and stained with an aqueous solution of DAPI (4′,6-diamidino-2-fenylindole). They were then analyzed with a Cyflow ® PA, Partec flow cytometer, taking as reference a sample with known diploid nuclei from the mango cultivar ‘Gomera3’ (Doležel at al., 1989; de Rocher *et al*., 1990).

DNA extraction was performed on young leaves of each of the four mango trees. Nine SSR were selected due to their high polymorphism for analysis of nuclear simple sequence repeats (Viruel *et al*., 2005). Each PCR reaction was performed using standard methodology, repeated at least twice to ensure reproducibility, and a range of mango accessions was used as positive controls to guarantee size accuracy and to minimize run-to-run variation.

### Leaf water content

To estimate leaf mass area (LMA) and water mass area (WMA), we first weighed 24 fresh leaves (six from each tree, or twelve per ploidy), then incubated in the oven at 60°C for 48h to get the dry weight (Poorter *et al*., 2009). We collected five additional leaves per tree, imaged them for leaf area (L_A_), rehydrated them at full turgor, and let them dry at room temperature, weighing them at regular intervals during one week until their weight was stable. We then obtained the dry weight, and calculated relative water loss on a per time basis (Turner, 1981).

The water potential of leaves (ψ_1_) and stems (ψ_s_) were measured weekly in the field from January to March, and from July to August 2020, using a Scholander pressure chamber (Scholander *et al*., 1965) model 600 (PMS Instrument, Corvallis, Oregon, USA). Six exposed leaves per tree (n=24 leaves per day of measurement) at similar phenological stages were evaluated. For ψ_s_, leaves were covered with a sealed foil bag 24h before measurements.

Twenty-eight additional leaves (12 from the diploid and 16 from the tetraploid trees), were used for the calculation of pressure/volume (P/V) curves in the laboratory (Sack *et al*., 2003). The petioles of the leaves were immersed in distilled water for 2h prior to obtaining the initial weight with a four-digit precision scale, then leaf water potential was measured and samples weighted again at regular intervals of 10 min (average temperature 19.5±0.05 □C and 52±0.39 % humidity), and this cycle was repeated at longer intervals until there was no more measurable water loss. Finally, leaves were dried in the oven at 60 °C for 48h, and weighed. These data served to infer the parameters of elasticity modulus (□), osmotic potential at full turgor (Π_0_), osmotic potential at turgor loss point Π_TLP_), relative water content at turgor loss point (RWC_TLP_), relative capacitance at full turgor (C’_FT_), and the symplastic/apoplastic fraction of water (s_f_, a_f_) (Koide *et al*., 1989).

### Anatomy of the leaves and immunolocalization of branched pectins

To compare the size of stomata between leaves from the diploid and tetraploid trees, we used peels of transparent nail polish applied to the abaxial leaf surface (Karabourniotis *et al*., 2001), which printed the guard cells and the thick cuticle, but not the profile of the epidermal cells. To get a better resolution of stomatal size, we fixed leaves in a solution containing acetic acid and methanol 1:1 (v/v) for 24h, stained them with the Feulgen reactive prior to clearing (Lora *et al*., 2017), and obtained 3D images of the stomata with a Leica confocal microscope (Leica Microsystems, Wetzlar, Germany).

The other half of the fixed leaves were washed with distilled water, dehydrated in a graded series of acetone, prior to incubation in Technovit 8100 resin for several days, and finally hardened under anoxic conditions at 4 °C. Sections of 4μm thick were obtained with a Leica EM UC7 ultramicrotome (Leica Microsystems, Wetzlar, GE). A monoclonal antibody that detects a branched α-galacturonan of some wall-related pectins (LM26, Plant Probes, Leeds, UK) was used. The sections were immunolocalized according to Torode *et al*., 2018, and counterstained with calcofluor white. Negative controls were treated in the same way, but substituting the primary antibody by a solution of 1% BSA in PBS.

### Geometry of the elements composing the xylem and the phloem in leaves, stems, and inflorescences

To evaluate the size of the vascular elements we cut small windows containing xylem and phloem tissues, placed them in a solution of hydrogen peroxide: acetic acid 1:1 (v/v), and incubated at 60 °C for at least two days (Franklin, 1945). Following tissue digestion, the debris was mounted onto glass slides with a drop of glycerol. To obtain phloem conduits, the same areas were longitudinally sectioned (i.e. parallel to the veins) in 1X Phosphate Buffer Saline (PBS) with a micro scalpel, to obtain ca. 1mm sections that were stained onto a microscope slide with a combination of 0.1% aniline blue in PO_4_ K_3_ (Currier and Strugger, 1956), which stains callose that accumulates in the sieve pore connections between the sieve tubes, and 0.1% calcofluor white in 10mM CHES buffer with 100mM KCl (pH=10) (Hughes and McCully, 1975) for cellulose. All slides were observed under a Leica DM LB2 upright microscope equipped with a Leica DM Camera and the LAS software, using differential interface contrast for vessel elements, and epifluorescence (405nm filter) for sieve tubes. We then measured a minimum of 50 conductive elements per leaf vein, per branch diameter, and per inflorescence location (n=1200 vessels and 1200 sieve tube elements). Finally, in order to refer our hydraulic calculations to the length covered by each conduit class, we measured the length of each sampling area (five per area) in each tree, and used the average to offer a picture of the real distance.

The radius of the simple perforation plates of the xylem were directly measured from the images obtained, but measurements of the sieve plate pore size of the phloem required further processing. Three samples from leaf veins (except from the primary vein), stem diameters, and inflorescence locations were obtained from trees of the two ploidies, immediately frozen in liquid nitrogen, kept in super chilled ethanol at −80° overnight, and gradually transferred to increasing temperatures of −20 °C, 4 °C, and room temperature (Mullendore *et al*., 2010). They were then cut at 2mm thickness with a double-edged razor blade, incubated in a 60 °C bath within a solution of 0.1% p/v proteinase K dissolved in 50 mM Tris-HCl buffer, and 8% Triton X-100 (pH 8,0), for two weeks. After that, they were dried with a freeze point dryer (CoolSafe 4–15L Freeze Dryers, LaboGene, Allerod, Denmark), mounted, and coated with gold palladium in a sputter coater (QUORUM Q 150 R ES), prior to observation with a LEICA scanning electron microscope (JEOL JSM-840). A total number of 1440 pores were analyzed. All microscopic images were processed with the Image J software (National Institutes of Health, Bethesda, MD).

### Hydraulic model

The anatomical data of xylem and phloem were used to model the resistance of the elements across the transport pathway, assuming a continuum between the stems and the different vein orders of leaves, or between the stem and the upright inflorescences. The mean vessel element radii and the mean sieve tube conduit radii were used for comparison. Based on previous reports (Knoblauch *et al*., 2016; Savage *et al*., 2017), we calculated the resistances applying the Hagen-Poiseuille model of laminar transport, which accounts for pressure driven flow in cylindrical tubes without axial transport. To compute the hydraulic resistance of the two vascular tissues, we followed the equation R= 8□l/πr^4^, with R being resistance (Pa s m^−3^), □ the viscosity of the fluid (mPa s), l the length of the conducting cell (m), and r the radius of the conducting cell (m). We assumed a standard constant viscosity of 1.0mPa s for the water within the xylem, and 1.7 mPa s for the sap within the phloem (Jensen *et al*., 2012). The total resistance of the conduits is the sum of the lumen resistance plus the resistance offered by the connections between tubes, as to R_tube_ = R_lumen_ + R_plate_. Given that vessel lumen resistances accounts for, at least, 50% of the total resistance of the vessel tube, and to compare both the xylem and the phloem tubes, we calculated the lumen resistances of both conducting elements for each defined class (for a detailed explanation of how R_lumen_ was calculated, see Knoblauch *et al*., 2016). To standardize these measurements with distance, we multiplied the R_lumen_ of each class by the adimensional factor L/l, being L 1m distance, and l the length of the conduit, which offers an idea of the number of individual conduits in one meter of the transport pathway, and thus, the resistivity of the continuous tubes. While the R_lumen_ of the sieve tube offers a good comparative framework between vascular tissues, in reality, the sieve plates contribute enormously to the total resistance of the sieve tubes. Therefore, their resistance was calculated, following the models developed by Jensen *et al*., 2012, and incorporated to the total resistance of the sieve tube in each class (see Savage *et al*., 2017; Clerx *et al*., 2020).

### Statistical analysis

All data were tested for normality with the Kolmogorov-Smirnov test (or the Saphiro Wilk test when n<50), and, when needed, log-transformed to fit normality. Physiological and anatomical parameters were compared within and between ploidies. Anatomical variation within the same ploidy was tested among sampling areas, using ANOVA and the Tukey test. For variation between ploidies, pairwise comparisons were tested with the t-test for similar sampling locations. Regression analysis were further applied to p/v curves to estimate the turgor loss point, and to the scaling of vascular element size. Stomatal lengths, density, and leaf areas were normalized to their maximum values for comparison. Data were analyzed with the SPSS software (SPSS Inc., Chicago, USA).

## Acknowledgements

We thank the laboratory of Professor José (Pepe) Escalona, in particular Ignacio Tortosa, from the University of Illes Balears, for their help with the PV curves. We also thank Sonia Cívico, Librada Alcaraz, and Yolanda Verdún, for their help with molecular and flow cytometry analysis. Laura Carrera very kindly provided the artwork. MB was financed by a JAE-ICU scholarship from the CSIC. JML was financed by a ComFuturo Project from the FGCSIC, and by a RTI Project (100-900-0000/AEI/10.13039/501100011033) from the Agencia Estatal de Investigación-Ministerio de Ciencia e Innovación, Spain, and LINKB20067 from CSIC. JIH was financed by a P18-RT-3272 project from Junta de Andalucía and by the Agencia Estatal de Investigación-Ministerio de Ciencia e Innovación, Spain (PID2019-109566RB-I00/AEI/10.13039/501100011033).

## Author contribution

M.B.A. performed field and laboratory work, and processed all data. N.M.H. co-designed, supervised the work, and wrote the manuscript. J.I.H. supervised the work and wrote the manuscript. J.M.L. designed the work, performed laboratory analysis, and wrote the manuscript.

## Data Availability Statement

The data that support the findings of this study are available from the corresponding author upon reasonable request.

## Supporting Information

**Fig. S1.** Scanning electron microscopy of the abaxial surface of a diploid mango leaf, showing the smooth continuous cuticle layer and the area of the stomatal guard cells corresponding with pores (black holes), where cuticle is lacking. Scale bar = 10μm.

**Fig. S2. Water potential of leaves (empty circles), and stems (solid circles) of diploid (pink) and tetraploid (green) mango ‘Kensington’.** From January to March, no differences were found in leaf or stem water potential, but during July and August, tetraploid mango trees reached slightly lower leaf water potentials. Bars represent standard error with *P* ≤ 0.05.

**Fig. S3.** Correlation between sieve area size and number of plates per end tube, showing a gradual increase in size and number from smaller to larger veins in leaves (open circles), and from thinner towards thicker branches (solid circles). Tetraploids (green) display higher number of plates and larger size compared with diploids (pink) at all locations along the transport pathway, including branches (filled circles), inflorescences (hatched circles), and leaves (empty circles). Vertical and horizontal bars represent standard errors at a *P* ≤ 0.05.

**Fig. S4.** Total number of pores per compound sieve plate of the phloem, comparing diploid (pink) and tetraploid (green) mango trees, along the different transport pathway. Bars represent the accumulated standard error at a *P* ≤ 0.05.

**Fig. S5. General morphology of mango trees from the reproductive phase (a,b) to initial fructification (c,d). (a)** The inflorescences display an upright position in the tree canopy during flowering. **(b)** Thousands of tiny male and hermaphrodite flowers form the mango inflorescences. **(c)** Most flowers abscise during the initial fruit set. **(d)** One to two fruits set per inflorescence, typically those located in the apical position (yellow arrow).

## REFERENCES

Barceló-Anguiano M, Hormaza JI, Losada JM. 2021. Conductivity of the phloem in mango (*Mangifera indica* L.) *Horticulture Research* (*in press*).

Bartlett MK, Scoffoni C, Ardy R, Zhang Y, Sun S, Cao K, Sack L. 2012. Rapid determination of comparative drought tolerance traits: using an osmometer to predict turgor loss point. Methods in Ecology and Evolution 3: 880–888.

Beaulieu JM, Leitch IJ, Patel S, Pendharkar A, Knight CA. 2008. Genome size is a strong predictor of cell size and stomatal density in angiosperms. New Phytologist 179: 975–986.

Bennett MD. 1972. Nuclear DNA content and minimum generation time in herbaceous plants. Proceedings of the Royal Society of London. Series B. Biological Sciences 181: 109–135.

Brodribb TJ, Feild TS. 2010. Leaf hydraulic evolution led a surge in leaf photosynthetic capacity during early angiosperm diversification. Ecology Letters 13: 175–183.

Brodribb TJ, Field TS, Sack L. 2010. Viewing leaf structure and evolution from a hydraulic perspective. Functional Plant Biology 37: 488–498.

Caine RS, Yin X, Sloan J, Harrison EL, Mohammed U, Fulton T, Biswal AK, Dionora J, Chater CC, Coe RA et al. 2018. Rice with reduced stomatal density conserves water and has improved drought tolerance under future climate conditions. New Phytologist 221: 371–384.

Carvalho MR, Turgeon R, Owens T, Niklas KJ. 2017. The scaling of the hydraulic architecture in poplar leaves. New Phytologist 214: 145–157.

Carvalho MR, Losada JM, Niklas KJ. 2018. Phloem networks in leaves. Current Plant Biology 43: 29–35.

Clerx LE, Rockwell FE, Savage JA, Holbrook NM. 2020. Ontogenetic scaling of phloem sieve tube anatomy and hydraulic resistance with tree height in *Quercus rubra*. American Journal of Botany 107: 852–863.

Comai L. 2005. The advantages and disadvantages of being polyploid. Nature Reviews Genetics 6: 836–846.

Currier HB, Strugger S. 1956. Aniline blue and fluorescence microscopy of callose in bulb scales of *Allium cepa* L. Protoplasma 45: 552–559.

De Baerdemaeker NJ, Hias N, Van den Bulcke J, Keulemans W, Steppe K. 2018. The effect of polyploidization on tree hydraulic functioning. American Journal of Botany 105: 161–171.

De Rocher EJ, Harkins KR, Galbraith DW, Bohnert HJ. 1990. Developmentally regulated systemic endopolyploid in succulents with small genomes. Science 250: 99–101.

Decanter L, Colling G, Elvinger N, Heiðmarsson S, Matthies D. 2020. Ecological niche differences between two polyploid cytotypes of *Saxifraga rosacea*. American Journal of Botany 107: 423–435.

Dixon HH, Joly J. 1895. The path of the transpiration-current. Annals of Botany 9: 403–420.

Doležel J, Binarová P, Lcretti S. 1989. Analysis of nuclear DNA content in plant cells by flow cytometry. Biologia Plantarum 31: 113–120.

Doyle JJ, Coate JE. 2019. Polyploidy, the nucleotype, and novelty: the impact of genome doubling on the biology of the cell. International Journal of Plant Sciences 180: 1–52.

Feild TS, Brodribb TJ. 2013. Hydraulic tuning of vein cell microstructure in the evolution of angiosperm venation networks. New Phytologist 199: 720–726.

Franklin GL. 1945. Preparation of thin sections of synthetic resins and wood-resin composites, and a new macerating method for wood. Nature 155: 51–51.

Ganino T, Rapoport HF, Fabbri A. 2011. Anatomy of the olive inflorescence axis at flowering and fruiting. Sciencia Horticulturae 129: 213–219.

García-García AL, Grajal-Martín MJ, González-Rodríguez Á. 2020. Polyploidization enhances photoprotection in the first stages of *Mangifera indica*. Scientia Horticulturae 264: 109198.

Gleason SM, Blackman CJ, Gleason ST, McCulloh KA, Ocheltree TW, Westoby M. 2018. Vessel scaling in evergreen angiosperm leaves conforms with Murray’s law and area filling assumptions: implications for plant size, leaf size and cold tolerance. New Phytologist 218: 1360–1370.

Hao GY, Lucero ME, Sanderson SC, Zacharias EH, Holbrook NM. 2013. Polyploidy enhances the occupation of heterogeneous environments through hydraulic related trade□offs in *Atriplex canescens* (Chenopodiaceae). New Phytologist 197: 970–978.

Harashima H, Schnittger A. 2010. The integration of cell division, growth and differentiation. Current Opinion in Plant Biology 13: 66–74.

Harrison EL, Arce Cubas L, Gray JE, Hepworth C. 2020. The influence of stomatal morphology and distribution on photosynthetic gas exchange. Plant Journal 101: 768–779.

Hughes J, Hepworth C, Dutton C, Dunn JA, Hunt L, Stephens J, Waugh R, Cameron DD, Gray JE. 2017. Reducing stomatal density in barley improves drought tolerance without impacting on yield. Plant Physiology 174: 776–787.

Hughes J, McCully ME. 1975. The use of an optical brightener in the study of plant structure. Stain Technology 50: 319–329.

Ietswaart R, Rosa S, Wu Z, Dean C, Howard M. 2017. Cell-size-dependent transcription of FLC and its antisense long non-coding RNA COOLAIR explain cell-to-cell expression variation. Cell Systems 4: 622–635.

Jensen KH, Mullendore DL, Holbrook NM, Bohr T, Knoblauch M, Bruus H. 2012. Modeling the hydrodynamics of phloem sieve plates. Frontiers Plant Science 3: 151.

Karabourniotis G, Tzobanoglou D, Nikolopoulos D, Liakopoulos G. 2001. Epicuticular phenolics over guard cells: exploitation for in situ stomatal counting by fluorescence microscopy and combined image analysis. Annals of Botany 87: 631–639.

Katagiri Y, Hasegawa J, Fujikura U, Hoshino R, Matsunaga S, Tsukaya, H. 2016. The coordination of ploidy and cell size differs between cell layers in leaves. Development 143: 1120–1125.

Knoblauch M, Knoblauch J, Mullendore DL, Savage JA, Babst BA, Beecher SD, Dodgen AD, Jensen KH, Holbrook NM. 2016. Testing the Münch hypothesis of long distance phloem transport in plants. Elife 5: e15341.

Koide RT, Robichaux RH, Morse SR, Smith CM, Pearcy RW, Ehleringer JR, Mooney HA, Rundel PW. 1989. Plant physiological ecology. Germany: Springer-Verlag, 161–183.

Kondorosi E, Roudier F, Gendreau E. 2000. Plant cell-size control: growing by ploidy? Current Plant Biology 3:488–492.

Levionnois S, Ziegler C, Jansen S, Calvet E, Coste S, Stahl C, Salmon C, Delzon S, Guichard C, Heuret P. 2020. Vulnerability and hydraulic segmentations at the stem–leaf transition: coordination across Neotropical trees. New Phytologist 228: 512–524.

Lewis WH. 1980. Polyploidy in species populations. New York: Plenum Press.

Li WD, Biswas DK, Xu H, Xu CQ, Wang XZ, Liu JK, Jiang G. 2009. Photosynthetic responses to chromosome doubling in relation to leaf anatomy in *Lonicera japonica* subjected to water stress. Functional Plant Biology 36: 783–792.

Li WL, Berlyn GP, Ashton PMS. 1996. Polyploids and their structural and physiological characteristics relative to water deficit in *Betula papyrifera* (Betulaceae). American Journal of Botany 83: 15–20.

Liesche J, Gao C, Binczycki P, Andersen SR, Rademaker H, Schulz A, Martens HJ. 2019. Direct comparison of leaf plasmodesma structure and function in relation to phloem-loading type. Plant Physiology 179: 1768–1778.

Liesche J, Pace MR, Xu Q, Li Y, Chen S. 2017. Height□related scaling of phloem anatomy and the evolution of sieve element end wall types in woody plants. New Phytologist 214: 245–256.

Liu H, Gleason SM, Hao G, Hua L, He P, Goldstein G, Ye Q. 2019. Hydraulic traits are coordinated with maximum plant height at the global scale. Science Advances 5: eaav1332.

Lora J, Herrero M, Tucker MR, Hormaza JI. 2017. The transition from somatic to germline identity shows conserved and specialized features during angiosperm evolution. New Phytologist 216:495–509.

Losada JM, Holbrook NM. 2019. Scaling of phloem hydraulic resistance in stems and leaves of the understory angiosperm shrub *Illicium parviflorum*. American Journal of Botany. 106: 244–259.

Lumaret R, Guillerm JL, Delay J, Lhaj Loutfi AA, Izco A, Jay M. 1987. Polyploidy and habitat differentiation in Dactylis glomerata L. from Galicia (Spain). Oecologia 73: 436–446.

Maherali H, Walden AE, Husband BC. 2009. Genome duplication and the evolution of physiological responses to water stress. New Phytologist 184: 721–731.

Masterson J. 1994. Stomatal size in fossil plants: evidence for polyploidy in majority of angiosperms. Science 264: 421–424.

Meyers LA, Levin DA. 2006. On the abundance of polyploids in flowering plants. Evolution 60: 1198–1206.

Mullendore DL, Windt CW, Van As H, Knoblauch M. 2010. Sieve tube geometry in relation to phloem flow. Plant Cell 22: 579–593.

Münch E. 1030. Die Stoffbewegungen in der Pflanze. Jena: Gustav Fischer.

Olson ME, Anfodillo T, Rosell JA, Petit G, Crivellaro A, Isnard S, León-Gómez C, Alvarado-Cárdenas LO, Castorena M. 2014. Universal hydraulics of the flowering plants: vessel diameter scales with stem length across angiosperm lineages, habits and climates. Ecology Letters 17: 988–997.

Olson ME, Soriano D, Rosell JA, Anfodillo T, Donoghue MJ, Edwards EJ, León-Gómez C, Dawson T, Camarero Martínez JJ, Castorena M et al. 2018. Plant height and hydraulic vulnerability to drought and cold. Proceedings of the National Academy of Sciences USA 115: 7551–7556.

Pei Y, Yao N, He L, Deng D, Li W, Zhang W. 2019. Comparative study of the morphological, physiological and molecular characteristics between diploid and tetraploid radish (*Raphunas sativus* L.). Sciencia Horticulurae 257: 108739.

Pickard WF. 2007. Laticifers and secretory ducts: two other tube systems in plants. New Phytologist 177: 877–888.

Poorter H, Niinemets Ü, Poorter L, Wright IJ, Villar R. 2009. Causes and consequences of variation in leaf mass per area (LMA): a meta analysis. New Phytologist 182: 565–588.

Ray DM, Savage JA. 2020. Immunodetection of cell wall pectin galactan opens up new avenues for phloem research. Plant Physiology 183: 1435–1437.

Roddy AB, Théroux-Rancourt G, Abbo T, Benedetti JW, Brodersen CR, Castro M, Castro S, Gilbride AB, Jensen B, Jiang GF et al. 2020. The scaling of genome size and cell size limits maximum rates of photosynthesis with implications for ecological strategies. International Journal of Plant Sciences 181: 75–87.

Romero-Aranda R, Bondada BR, Syvertsen JP, Grosser JW. 1997. Leaf characteristics and net gas exchange of diploid and autotetraploid citrus. Annals of Botany 79: 153–160.

Ruiz M, Oustric J, Santini J, Morillon R. 2020. Synthetic polyploidy in grafted crops. Frontiers Plant Science 11: 540894.

Sack L, Holbrook NM. 2006. Leaf hydraulics. Annual Review Plant Biology 57: 361–381.

Sack L, Scoffoni C. 2013. Leaf venation: structure, function, development, evolution, ecology and applications in the past, present and future. New Phytologist 198: 983–1000.

Sack L, Cowan PD, Jaikumar N, Holbrook NM. 2003. The ‘hydrology’of leaves: co□ordination of structure and function in temperate woody species. Plant Cell & Environment 26: 1343–1356.

Galán-Saúco V, Grajal-Martín MJ, Fernández-Galván D, Coello-Torres A, Juárez J, Navarro L. 2001. Occurrence of spontaneous tetraploid nucellar mango plants. HortScience 36: 755–757.

Savage JA, Beecher SD, Clerx L, Gersony JT, Knoblauch J, Losada JM, Jensen KH, Knoblauch M, Holbrook, NM. 2017. Maintenance of carbohydrate transport in tall trees. Nature Plants 3: 965–972.

Savage JA. 2019. A temporal shift in resource allocation facilitates flowering before leaf out and spring vessel maturation in precocious species. American Journal of Botany 106: 113–122.

Savage JA, Chuine I. 2021. Coordination of spring vascular and organ phenology in deciduous angiosperms growing in seasonally cold climates. New Phytologist 230: 1700–1715.

Scholander PF, Bradstreet ED, Hemmingsen EA, Hammel HT. 1965. Sap pressure in vascular plants: negative hydrostatic pressure can be measured in plants. Science 148: 339–346.

Sevanto S. 2018. Drought impacts on phloem transport. Current Opinion in Plant Biology 43: 76–81.

Simonin KA, Roddy AB 2018. Genome downsizing, physiological novelty, and the global dominance of flowering plants. PLoS Biology 16: e2003706.

Smith HE. 1946. *Sedum pulchellum:* a physiological and morphological comparison of diploid, tetraploid, and hexaploid races. Journal of the Torrey Botanical Society 73: 495–541.

Snodgrass SJ, Jareczek J, Wendel JF. 2017. An examination of nucleotypic effects in diploid and polyploid cotton. AoB Plants 9.

Stebbins GL. 1950. Variation and evolution in plants. London, UK: Oxford University Press.

Tal M, Gardi I. 1976. Physiology of polyploid plants: water balance in autotetraploid and diploid tomato under low and high salinity. Physiologia Plantarum 38: 257–261.

Torode TA, O’Neill R, Marcus SE, Cornuault V, Pose S, Lauder RP, Kracun SK, Rydahl MG, Andersen MC, Willats WG et al. 2018. Branched pectic galactan in phloem-sieve-element cell walls: implications for cell mechanics. Plant Physiology 176: 1547–1558.

Trifiló P, Raimondo F, Savi T, Lo Gullo MA, Nardini A. 2016. The contribution of vascular and extra-vascular water pathways to drought-induced decline of leaf hydraulic conductance. Journal of Experimental Botany 67: 5029–5039.

Turner NC. 1981. Techniques and experimental approaches for the measurement of plant water status. Plant Soil 58:339–366.

Van de Peer Y, Ashman TL, Soltis PS, Soltis DE. 2020. Polyploidy: an evolutionary and ecological force in stressful times. The Plant Cell 33: 11–26.

Viruel MA, Escribano P, Barbieri M, Ferri M, Hormaza JI. 2005. Fingerprinting, embryo type and geographic differentiation in mango (*Mangifera indica* L., Anacardiaceae) with microsatellites. Molecular Breeding 15: 383–393.

Vyas P, Bisht MS, Miyazawa SI, Yano S, Noguchi K, Terashima I, Funayama-Noguchi. 2007. Effects of polyploidy on photosynthetic properties and anatomy in leaves of *Phlox drummondii*. Functional Plant Biology 34: 673–682.

Lihu W, Luo Z, Wang L, Deng W, Wei H, Liu P, Liu M. 2019. Morphological, cytological and nutritional changes of autotetraploid compared to its diploid counterpart in Chinese jujube (*Ziziphus jujuba* Mill.). Scientia Horticulturae 249: 263–270.

Warschefsky EJ, von Wettberg EJ. 2019. Population genomic analysis of mango (*Mangifera indica*) suggests a complex history of domestication. New Phytologist 222: 2023–2037.

Wei N, Cronn R, Liston A, Ashman TL. 2019. Functional trait divergence and trait plasticity confer polyploid advantage in heterogeneous environments. New Phytologist 221: 2286–2297.

Wei N, Du Z, Liston A, Ashman TL. 2020. Genome duplication effects on functional traits and fitness are genetic context and species dependent: studies of synthetic polyploid Fragaria. American Journal of Botany 107: 262–272.

Xiong D, Nadal M. 2019. Linking water relations and hydraulics with photosynthesis. The Plant Journal 101: 800–815.

Xu C, Zhang Y, Huang Z, Yao P, Li Y, Kang X. 2018. Impact of the leaf cut callus development stages of *Populus* on the tetraploid production rate by colchicine treatment. Journal of Plant Growth Regulation 37: 635–644.

Zhang F, Xue H, Lu X, Zhang B, Wang F, Ma Y, Zhang Z. 2015. Autotetraploidization enhances drought stress tolerance in two apple cultivars. Trees 29: 1773–1780.

Zhang XY, Wang XL, Wang XF, Xia GH, Pan QH, Fan RC, Wu FQ, Yu XC, Zhang DP. 2006. A shift of phloem unloading from symplasmic to apoplasmic pathway is involved in developmental onset of ripening in grape berry. Plant Physiology 142: 220–232.

Zimmermann MH. 1983. Xylem structure and the ascent of sap. Germany: Springer-Verlag.

