## Supplementary figures and images for "Changes In Ploidy Affect Vascular Allometry And Hydraulic Function In Trees"

### Supplemental Figure 1

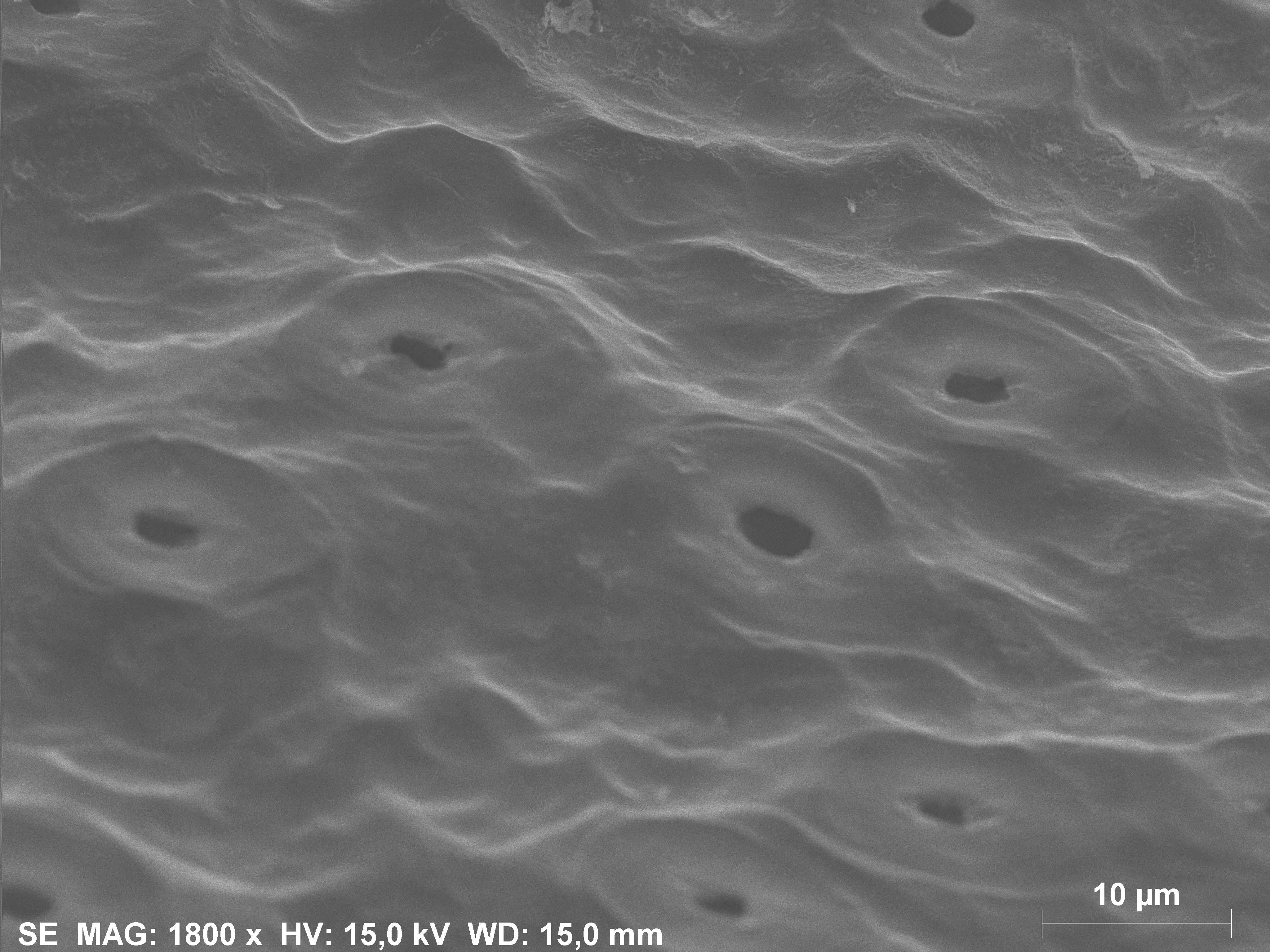

### Supplemental Figure 2

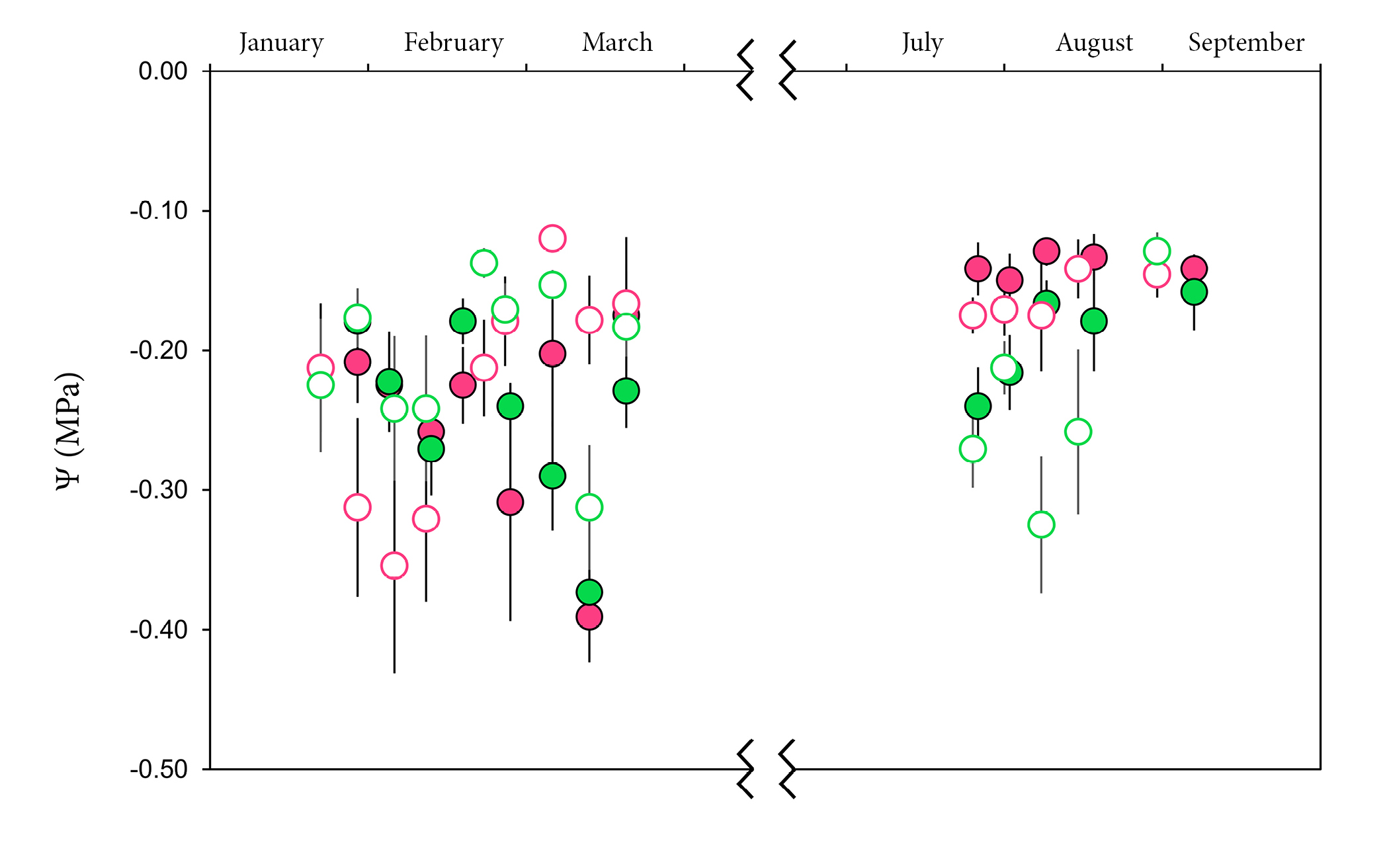

### Supplemental Figure 3

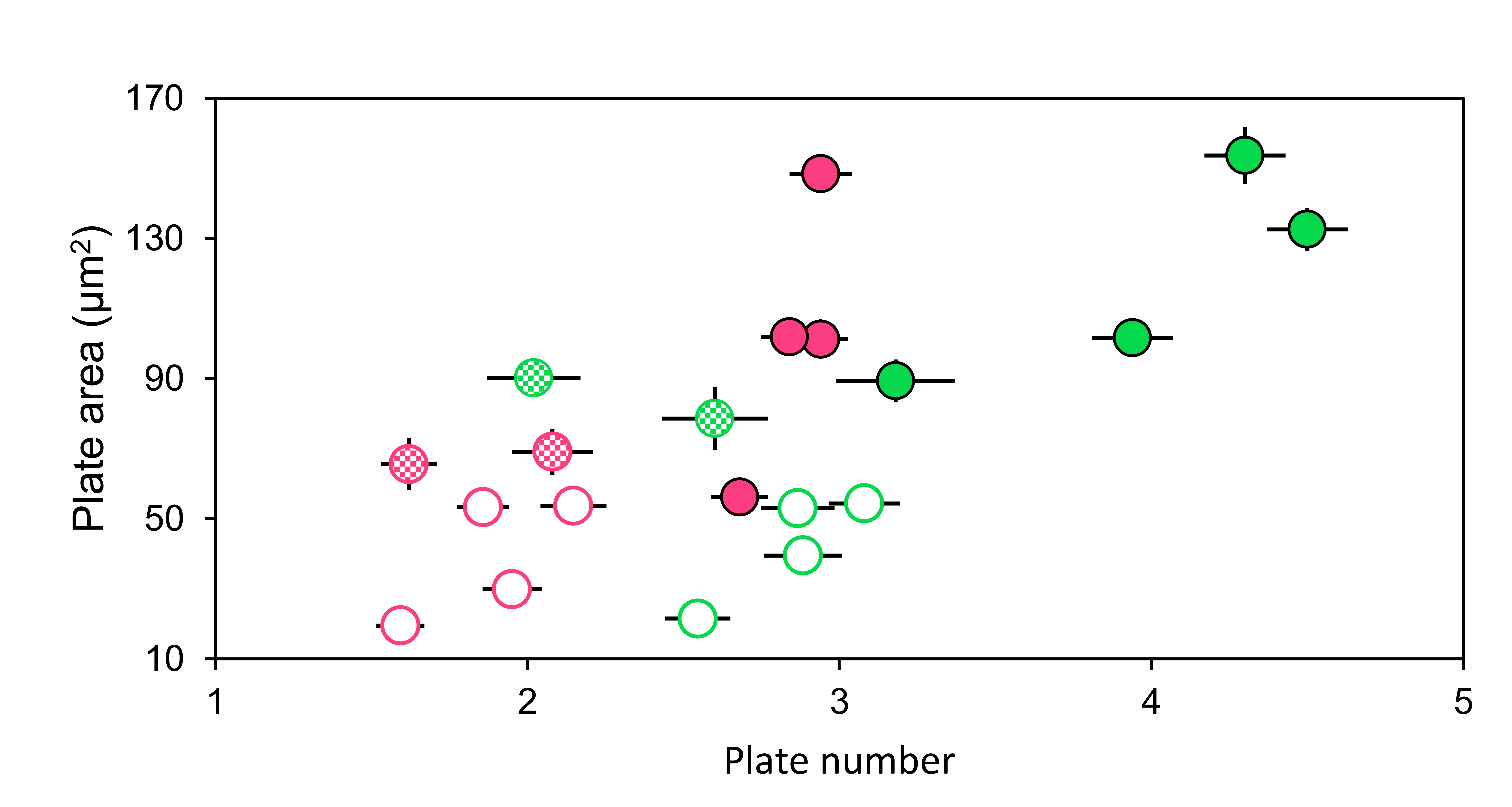

### Supplemental Figure 4

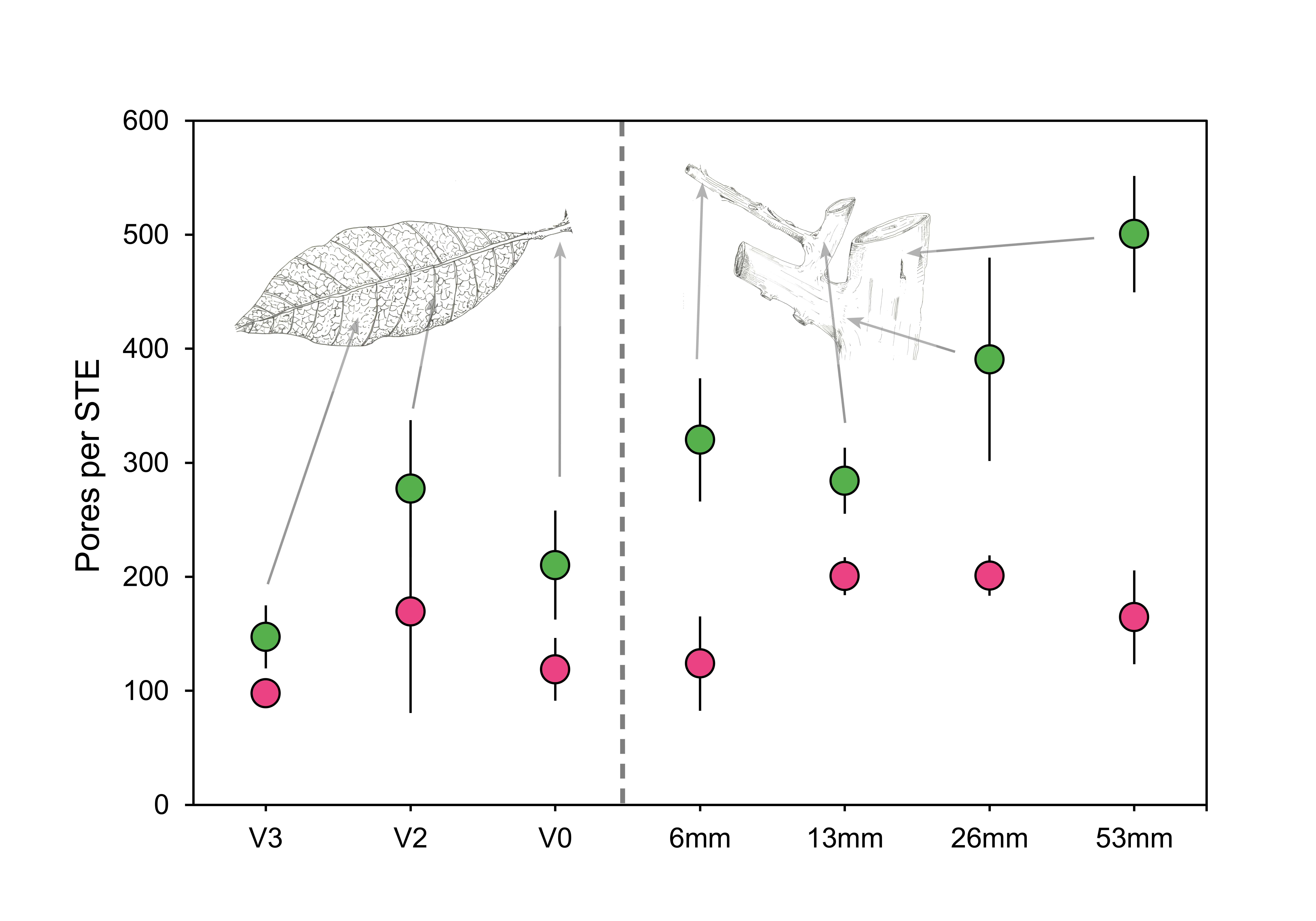

### Supplemental Figure 5

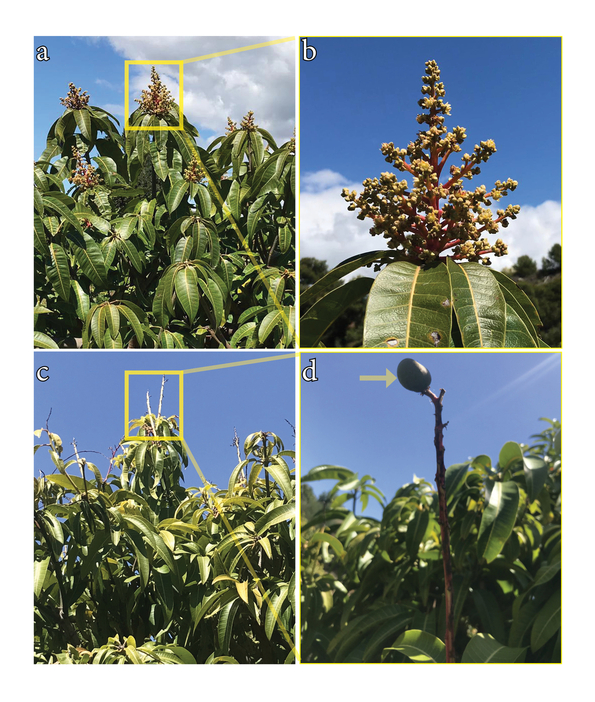
